# AMP-activated protein kinase (AMPK) is required for recovery from metabolic stress induced by ultrasound microbubble treatment

**DOI:** 10.1101/2022.03.02.482704

**Authors:** Louis Lo, Oro Uchenunu, Roberto J. Botelho, Costin N. Antonescu, Raffi Karshafian

**Affiliations:** Department of Chemistry and Biology, Ryerson University, Toronto, Ontario, Canada, M5B 2K3; Graduate Program in Molecular Science, Ryerson University, Toronto, Ontario, Canada, M5B 2K3; Faculty of Medicine and Health Sciences, Division of Experimental Medicine, McGill University, Montreal, Quebec, Canada, H3A 0G4; Department of Physics, Ryerson University, Toronto, Ontario, Canada, M5B 2K3; Institute for Biomedical Engineering, Science and Technology (iBEST), a partnership between Ryerson University and St. Michael’s Hospital, Toronto, Ontario, Canada, M5B 1W8; Keenan Research Centre for Biomedical Science of St. Michael’s Hospital, Toronto, Ontario, Canada, M5B 1W8

## Abstract

Ultrasound and microbubbles (USMB) is a promising strategy for cancer therapy. USMB can induce a variety of effects on cells including transient formation of plasma membrane pores (sonoporation) and enhanced endocytosis, which enhance drug delivery, and can also lead to enhanced cell death. However, the outcomes of USMB on cell physiology are heterogeneous, in that USMB elicits cell death in a proportion of cells while exerting minimal effects on others. This suggests that mechanisms of adaptation following USMB allow some cells to survive and/or proliferate. The molecular mechanisms of adaptation to USMB-induced stress remain poorly understood, thus potentially hindering broad therapeutic applications of USMB. Herein, we used several triple negative breast cancer cells to study the effect of USMB-induced metabolite stress and the role of AMPK as a response to this stress. We found that USMB alters steady-state levels of amino acids, glycolytic intermediates, and citric acid cycle intermediates. USMB treatment acutely reduces ATP levels and stimulates AMP-activated protein kinase (AMPK) phosphorylation and activation. Further, AMPK is required to restore ATP levels in cells that survived the initial insult and support cell proliferation post-USMB treatment. These results suggest that AMPK and metabolic perturbations are likely determinants of the anti-neoplastic efficacy of USMB treatment.

## Introduction

The application of ultrasound waves on biological tissues can induce a range of changes to cell and tissue physiology, and in the presence of microbubbles (MBs) - micron-sized, gas-filled spherical structures stabilized by a shell (lipid, protein, polymer) - these biological effects of ultrasound are amplified (Guzmán et al., 2003; Martin and Dayton, 2013). Combinations of MBs and ultrasound are being investigated for clinical applications including those aiming to improve targeted drug delivery in cancer (Goertz et al., 2012; Meijering et al., 2009; Sorace et al., 2012) and targeted delivery through the blood-brain-barrier (Dasgupta et al., 2016; Konofagou et al., 2012). The effectiveness of ultrasound-stimulated microbubble (USMB) treatment as a targeted delivery strategy in cell culture models was demonstrated for various agents including chemotherapy, gold nanoparticles, genetic material, and antibiotics (Huang et al., 2017; Sirsi and Borden, 2012; Sorace et al., 2012; Tarapacki and Karshafian, 2015). The effects of USMB are in large part mediated by enhanced uptake of extracellular material through pore formation on the plasma membrane, referred to as sonoporation (Helfield et al., 2016; Lentacker et al., 2014), and enhancement of endocytic uptake by several pathways (Derieppe et al., 2015; Fekri et al., 2019, 2016; Meijering et al., 2009). However, cells exposed to USMB alone may exhibit a variety of cell biological effects including activation of diverse signaling cascades which regulate cell survival, apoptosis, and cellular adaptation (Haugse et al., 2020, 2019; Zhang et al., 2012). As such, understanding the mechanisms that cells engage to adapt to USMB-induced cell stress is critical.

A less appreciated outcome of sonoporation is that this phenomenon also enhances the diffusion of cellular material to the extracellular milieu (Hussein et al., 2017). Consistent with this, we previously reported the loss of molecules such as cytosolic GFP-tagged proteins from the cell following USMB treatment (Hussein et al., 2017). This suggests that a range of intracellular molecules may also be released during sonoporation, leading to their depletion from within the cell (Hussein et al., 2017). We thus hypothesized that USMB may lead to the loss of key metabolites including nucleotides, glucose, glycolytic intermediates, and amino acids, and that loss of these may result in metabolic and signaling perturbations that can impact cell fate (Herzig and Shaw, 2018; Lin and Hardie, 2018; Yang and Vousden, 2016).

Cells respond to changes in available nutrients (Balakrishnan et al., 2018; Lanning et al., 2017; Lin and Hardie, 2018; Saxton and Sabatini, 2017; Yang et al., 2013; Yang and Vousden, 2016) by adjusting their metabolic pathways to maintain homeostasis (Cairns et al., 2011; Moon et al., 2019). Under periods of high nutrient availability, cellular signaling pathways enable anabolism and cell growth (Yang et al., 2013). In contrast, under periods of nutrient insufficiency, key biochemical pathways enable catabolism and restoration of metabolic homeostasis (Balakrishnan et al., 2018). For instance, AMP-activated protein kinase (AMPK) is a key cellular sensor of metabolic stress (Herzig and Shaw, 2018; Lin and Hardie, 2018). AMPK consist of a catalytic α subunit, and regulatory β and γ subunits (Oakhill et al., 2009). Cells experiencing nutrient depletion also often exhibit an increase in ADP and AMP levels relative to ATP. AMP/ADP competes for ATP to bind to some cystathionine β-synthetase domains on the γ subunits of AMPK, eliciting a conformational change that leads to enhanced phosphorylation of Thr172 on the α subunit by LKB1, leading to AMPK activation (Oakhill et al., 2009). Nutrient stress in the form of reduced glucose availability also activates AMPK in a manner independent of ATP levels (Zhang et al., 2017). AMPK is also activated in response to increases in intracellular calcium and mechanical stress transduced by E-cadherin, both requiring AMPK phosphorylation by LKB1 or CaMKK (Bays et al., 2017; Lin and Hardie, 2018; Oakhill et al., 2009; Shackelford and Shaw, 2009). Once activated, AMPK restores cellular metabolic balance, in general by decreasing anabolic processes, while stimulating catabolic pathways. For instance, AMPK decreases lipogenesis while stimulating fatty acid oxidation via phosphorylation and inactivation of acetyl-CoA-carboxylase (ACC) (Hardie and Pan, 2002). Moreover, AMPK can inactivate mTORC1 signaling thus resulting in the reduction of protein synthesis (Lin and Hardie, 2018).

In this work, we postulated that USMB triggers metabolic stress and activates AMPK, and that AMPK may consequently reprogram cells to support viability and/or proliferation of a subset of cells following USMB. To test this, we employed USMB on MDA-MB-231, SUM149PT, and BT-20 triple negative breast cancer cell lines. We used several approaches to reveal that USMB treatment leads to decreased intracellular levels of various metabolites while concurrently stimulating AMPK. We then found that AMPK is necessary for cells to recover from this USMB-mediated metabolic insult and proliferate following USMB treatment. This suggests that co-administration of AMPK inhibitors during USMB may be an effective anti-tumour strategy.

## Results

Considering that our previous study showed that USMB treatment causes efflux of GFP-tagged proteins from cells (Hussein et al., 2017), we set out to investigate the effects of USMB on the steady-state metabolome of surviving cells. To achieve this, we applied USMB on MDA-MB-231 breast cancer cells using a custom-built ultrasound transducer system (**Figure 1**). To establish the optimal conditions for USMB treatment of MDA-MB-231 cells, we monitored the influx of plasma-membrane impermeable dye, Lucifer Yellow by confocal microscopy (**Figure 2A**). USMB exposure increased the average fluorescence intensity of Lucifer Yellow per cell. We observed ~80% of USMB treated cells with Lucifer Yellow intensity (median=3.267) higher than the upper 95% confidence interval of the mean of the untreated cells (Basal, Upper 95% CI of mean=1.1039) (**Figure 2A**, right), indicating that this USMB treatment induced sonoporation. As a positive control, we perforated cell membrane with digitonin which resulted in significant increase in fluorescence intensity compared to untreated cells (**Figure 2A**).

**Figure 1.**
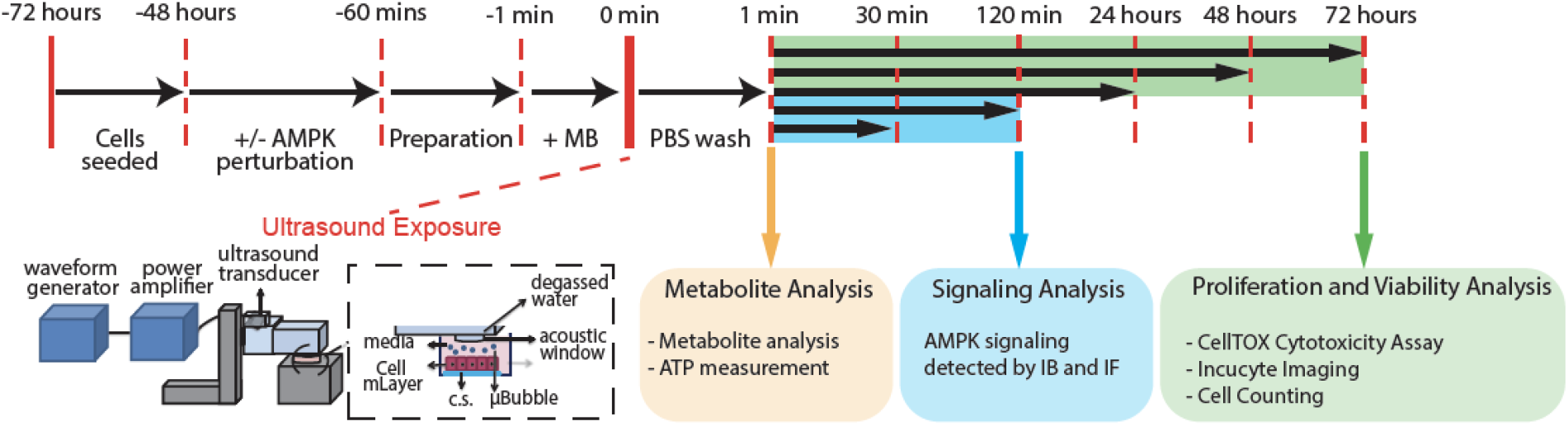
Design of the Study. MDA-MB-231, SUM149PT or BT-20 cells were prepared and treated with USMB using a calibrated acoustic exposure platform that was developed to deliver ultrasound pulses, as was previously described (Fekri et al., 2019). See Supplemental Information for additional details of USMB treatment. Numerous assays were performed following USMB treatment over a 72 h period as indicated, including metabolite analysis (Figure 2,4), Signaling Analysis (Figure 3,5) and proliferation and viability assays (Figure 6).

**Figure 2.**
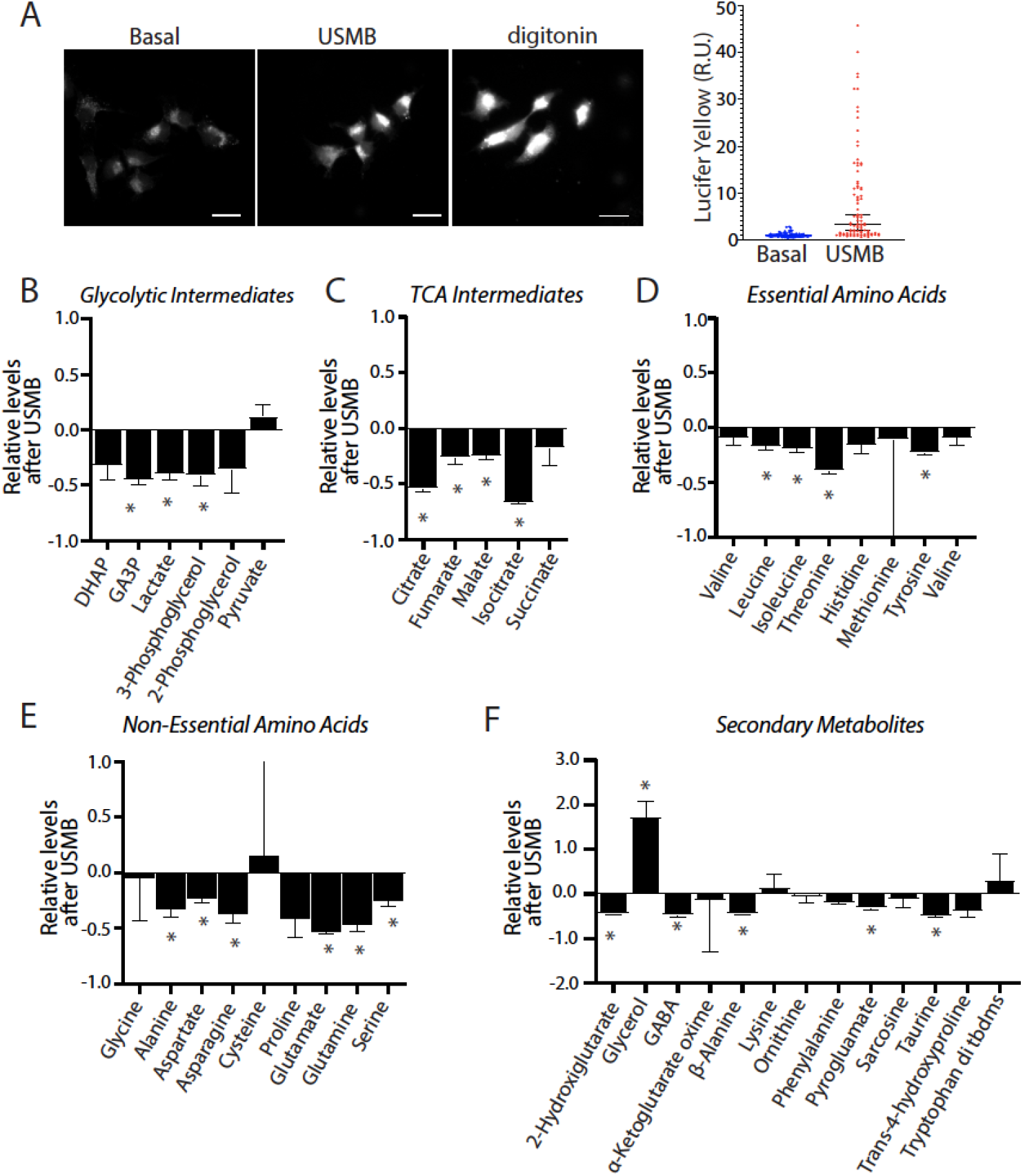
USMB impacts a broad range of cellular levels of metabolites and amino acids. (A) MDA-MB-231 cells in the presence of Lucifer Yellow Potassium Salt (125 μg/mL) were subject to USMB treatment (USMB) or 2μg/mL digitonin for 5 min, as indicated. Shown (left panels) are representative images obtained by widefield epifluorescence microscopy, scale bar = 100 μm. Also shown (right panels) is the quantification of Lucifer Yellow fluorescence, depicting fluorescence measured in individual cells (individual points) and median (horizontal bar) along with 95% confidence interval (CI). (B-F) Levels of indicated metabolites in MDA-MB-231 cells treated with USMB were determined by gas chromatography-mass spectrometry (GC/MS). Data are shown as a mean ± SD (n=3) expressed as levels detected in USMB samples relative to untreated control; Relative levels = ([USMB-treated]/[control]) - [control] for each metabolite. Statistical analysis was performed by unpaired t-tests (comparing control vs USMB-treated samples) for each metabolite, *, p< 0.05.

We hypothesized that USMB causes a loss of cytosolic contents such as glycolytic and citric acid cycle (TCA) metabolites and amino acids, which are directly involved in numerous aspects of cellular function and survival (Saxton and Sabatini, 2017). In order to assess this, intracellular steady-state levels of select metabolites were quantified post-USMB treatment. USMB reduced the levels of many glycolytic and citric acid cycle intermediates as well as essential and non-essential amino acids (**Figure 2B–F**). Notably, the extent of loss of various metabolites upon USMB varied, suggesting that following initial depletion of certain metabolites from cells after USMB treatment, cataplerosis and/or anaplerosis may partly replenish some metabolites at the time of the assay. For example, while USMB caused a significant decrease in glyceraldehyde-3-phosphate and 3-phosphoglycerol, the impact of USMB on DHAP and 2-phosphoglycerol appeared to be less pronounced. Similarly, the TCA intermediates citrate, fumarate, and malate were significantly reduced in USMB-treated cells, while changes in other TCA metabolites such as alpha-ketoglutarate and succinate were not observed following USMB (**Figure 2C**).

The intracellular levels of essential amino acids (leucine, isoleucine, and threonine), non-essential amino acids derived from citric acid cycle intermediates (glutamine, glutamate, and aspartate), and other non-essential amino acids (alanine, asparagine, tyrosine, and serine), were all also significantly reduced by USMB treatment (**Figure 2D–E**). As observed for some glycolytic and TCA intermediates, not all amino acids levels were uniformly affected by USMB treatment, as the levels of glycine, cysteine and proline did not exhibit appreciable changes following USMB treatment (**Figure 2D–E**). A number of secondary metabolites (2-hydroxyglutarate, GABA, β-alanine, ornithine, phenylamine, and taurine) were also significantly reduced relative to control, untreated cells (**Figure 2F**). We also observed a large accumulation of glycerol following USMB; however, this is likely due to the remnants of microbubble debris where glycerol is a key ingredient of the DEFINITY microbubble shell composition (Lantheus Medical Imaging, DEFINITY, MSDS, 10/4/2015). Overall, these results show that USMB-induced sonoporation of MDA-MB-231 cells has a broad effect to significantly reduce the intracellular levels of a diverse set of metabolites. This, in turn suggests that cells exposed to USMB treatment may suffer significant energetic stress due to loss of key metabolites and nutrients.

Given our observation that USMB reduced glycolytic intermediates and amino acids in surviving cells, we hypothesized that USMB-induced sonoporation induces AMPK activation. To determine whether USMB triggered AMPK activation, we probed for phosphorylation of AMPK (pT172-AMPK) and acetyl CoA-carboxylase (pS79-ACC), a substrate of AMPK, in three triple negative breast cancer cell models. Western blotting of whole cell lysates revealed that USMB treatment resulted in greater phosphorylation of pT172-AMPK and pS79-ACC in MDA-MB-231, SUM149PT, and BT-20 cells (**Figure 3A–C**). We complemented this approach with immunofluorescence microscopy to detect pS79-ACC upon USMB treatment in single MDA-MB-231 cells (**Figure 3D**). We found that pS79-ACC was increased following USMB treatment in comparison to the untreated control (**Figure 3E**). Furthermore, pharmacological inhibition of AMPK using 10 μM Compound C for 1 h prior to USMB reduced pS79-ACC fluorescence, indicating that the increase in ACC phosphorylation in USMB-treated conditions indeed reflected enhanced AMPK activation (**Figure 3E**).

**Figure 3.**
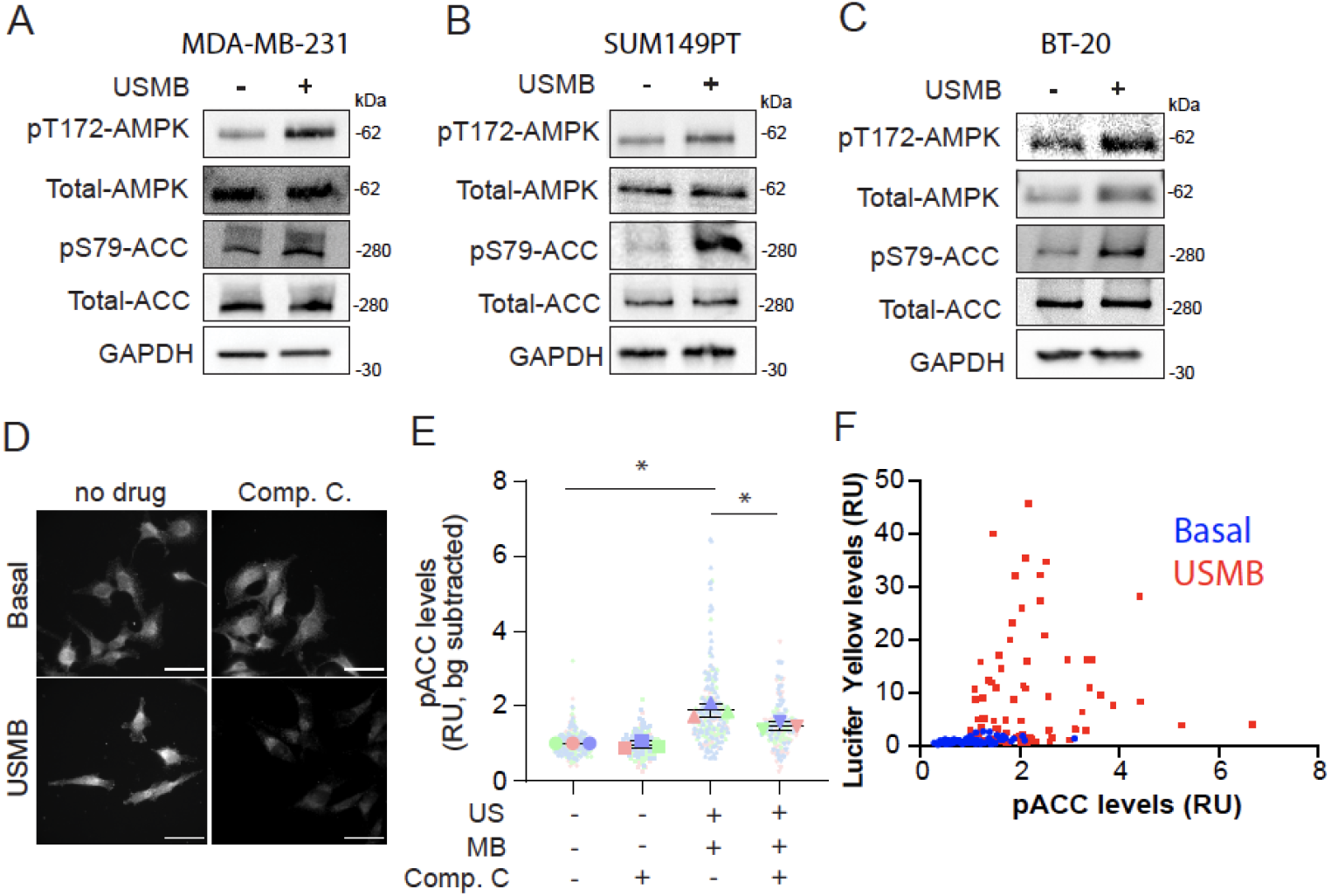
USMB triggers AMPK activation. **(A-C)** MDA-MB-231, SUM149PT, and BT-20 cells were treated with USMB or left untreated (basal). Whole cell lysates were prepared 30 min after USMB treatment and were analyzed by Western blotting with depicted antibodies. (D-E) MDA-MB-231 cells were fixed, permeabilized, and stained for endogenous pS79-ACC, 30 min after USMB treatment. Some samples were also pre-treated with 10 μM compound C for 1 h prior to USMB treatment, as indicated. Shown in (D) are representative images obtained by widefield epifluorescence microscopy, scale bar, 100μm. Shown in (E) is the quantification of immunofluorescence intensity; data is represented as normalized fluorescence (relative units, R.U.) values in individual cells (small circles), average cell fluorescence intensity in each experiment in each condition (large shapes) and mean of independent experiments (bar) ± SD (whiskers). The experiment was repeated three times independently. Measurements are color-coded by independent experiment. Statistical analysis was performed by one-way ANOVA followed by Tukey’s multiple comparisons test. *, p < 0.05. (F) A similar experiment was performed in which MDA-MB-231 were subject to USMB in the presence of Lucifer Yellow Potassium Salt (125μg/mL), then subject to fixation and staining. Shown is the quantification of immunofluorescence intensity of phospho-ACC and Lucifer Yellow for individual MDA-MB-231 cells treated with USMB as indicated.

The broad distribution of pS79-ACC staining intensity in cells upon USMB treatment suggests a heterogenous response to USMB treatment (**Figure 3E**). If the heterogeneity in AMPK activation following USMB reflects a heterogeneity in the extent of sonoporation, and thus the extent of loss of cellular metabolites, then the levels of sonoporation and AMPK activation should correlate in individual cells. To this end, we examined both pS79-ACC staining intensity with the uptake of Lucifer Yellow (a marker of sonoporation, Figure 2A) in response to USMB. The majority of USMB-treated cells that exhibited elevated pS79-ACC levels also had higher Lucifer Yellow fluorescence intensity (mean=8.190) (**Figure 3F**), indicating that sonoporation and AMPK activation co-occur.

Since we observed a general decrease across measured metabolites following USMB treatment, we hypothesized that AMPK activation plays a key role in restoring energy and metabolite balance following USMB-induced sonoporation. To test this, we measured ATP levels in the cells before and after USMB treatment in MDA-MB-231 cells. This revealed that USMB induces an acute loss of cytosolic ATP (after <5 min of USMB treatment, **Figure 4**). Importantly, cytosolic ATP levels were restored 1 h after USMB treatment (**Figure 4**, blue bars). To determine if AMPK plays a role in recovery of ATP levels following USMB treatment, we used short interfering RNA (siRNA) to silence the catalytic α subunit of AMPK (both α1 and α2 isoforms) in parallel experiments and quantified intracellular ATP levels following USMB treatment. In AMPK-depleted cells, we observed similar reduction in intracellular ATP levels immediately following USMB treatment as in control siRNA treated cells. However, in contrast to control siRNA treated cells, there was no appreciable recovery of ATP levels for up to 2 h post-treatment (**Figure 4**, red bars). These observations imply that USMB triggers a reduction in intracellular ATP levels, while AMPK is required for recovery of ATP levels.

**Figure 4.**
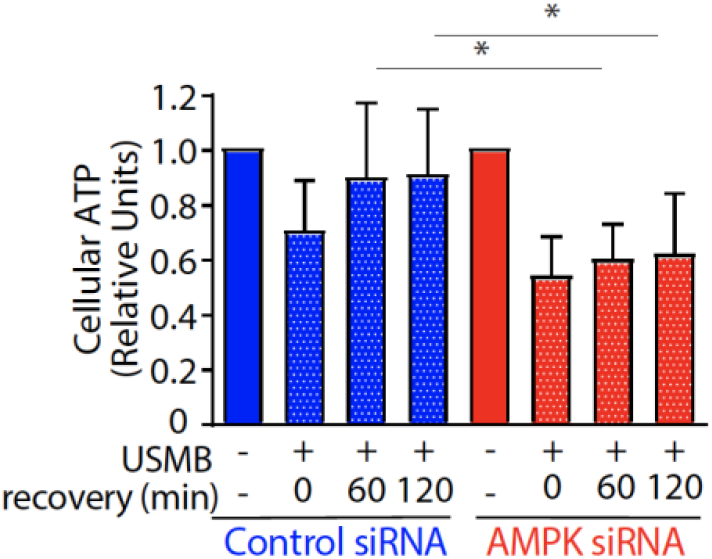
Recovery of cellular ATP levels following USMB treatment requires AMPK. MDA-MB-231 cells were treated with control (non-targeting) or AMPK siRNA. Following transfection, cells were treated with USMB as indicated, which was then followed by a recovery period as shown. Cell samples were then subject to a luminescence-based assay to measure ATP levels. ATP measurements were normalized to total protein and also normalized to the control (no USMB) condition. Shown are the mean ATP values ± SD, (n=7). Statistical analysis was performed by one-way ANOVA followed by Tukey’s multiple comparisons test. *, p < 0.05.

To study the effects of USMB and the role of AMPK on cell survival and growth following USMB treatment, we generated MDA-MB-231 and SUM149PT cells with doxycycline-inducible expression of short hairpin RNA (shRNA) targeting AMPK (MDA-shAMPK and SUM149PT-shAMPK henceforth). As expected, doxycycline treatment for 48 hours resulted in dose-dependent silencing of AMPKα1/2 in MDA-shAMPK cells (**Figure 5A**). Treatment of MDA-shAMPK cells with 3 μM doxycycline for 48 hours resulted in approximately 50-80% loss of the AMPKα1/2 subunit in MDA-shAMPK (**Figure 5B**) and SUM149PT-shAMPK (**Figure 5C**) cells. Importantly, in MDA-shAMPK, USMB-treated cells depleted of AMPK by doxycycline treatment exhibited reduced pT172-AMPK and pS79-ACC compared to cells treated with USMB and expressing AMPK (without doxycycline treatment; **Figure 5B**). Collectively, these results demonstrate that AMPK levels and/or activity was effectively suppressed in MDA-shAMPK and SUM149PT-shAMPK cells following addition of doxycycline.

**Figure 5.**
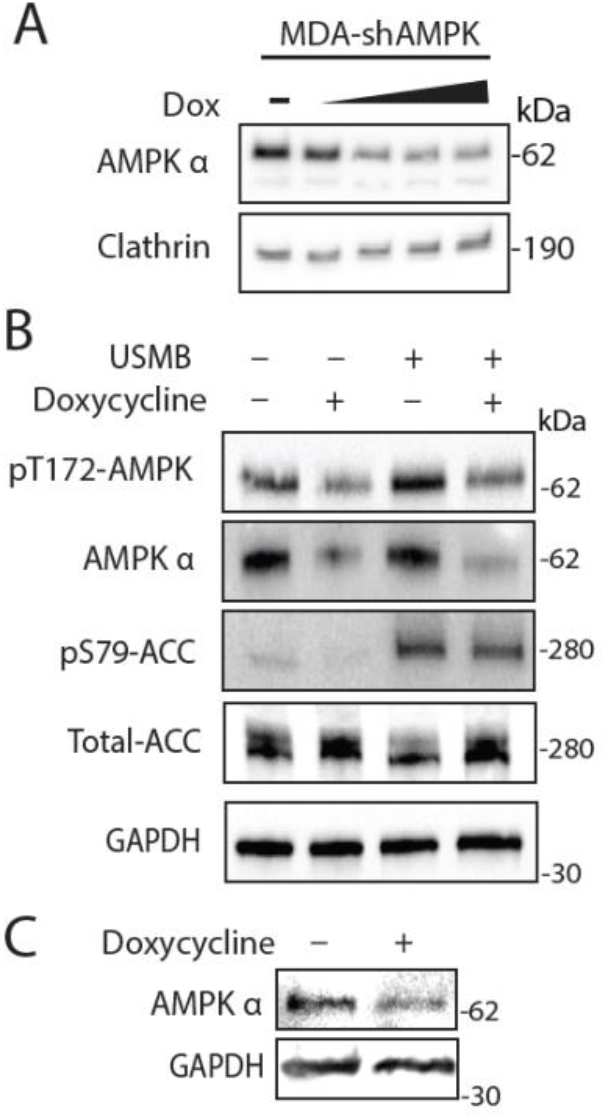
Inducible expression of AMPK shRNA achieves effective silencing of AMPK activity. (A) Expression of AMPK shRNA in MDA-shAMPK was induced by incubation of 0.1-5 μM doxycycline for 48 h. Shown are representative western blots of whole cell lysates probed with antibodies specific for AMPK α1/2 or clathrin (loading control). (B) Expression of AMPK shRNA in MDA-shAMPK was induced by incubation of 3 μM doxycycline for 48 h, as indicated. Cells were then also then treated with USMB, as indicated, and whole cell lysates were prepared 30 min after USMB treatment. Shown are representative western blots with antibodies as indicated. (C) Expression of AMPK in SUM149PT-shAMPK cells following incubation of 3 μM doxycycline for 48 h, as indicated. Shown are representative western blots with antibodies as indicated.

So far, our observations show that USMB elicits a reduction in the levels of many metabolites, including ATP. Cells that survive USMB recover ATP levels by 2 h post-treatment in an AMPK-dependent manner. We thus hypothesized that suppressing AMPK would lead to a decrease in cell viability and/or arrest cell proliferation in response to USMB. By counting the number of viable cells, we found that in the absence of USMB treatment, MDA-shAMPK cells treated with doxycycline to deplete AMPK exhibited comparable proliferation rates as control cells not treated with doxycycline (**Figure 6A**, blue and red lines). In contrast, AMPK depletion by doxycycline treatment led to significant reduction of the number of viable cells following USMB treatment (**Figure 6A**, orange line) relative to cells treated with USMB alone (without doxycycline treatment) (**Figure 6A**, green line). This suggests that AMPK is essential to support cell proliferation and/or viability of MDA-MB-231 cells upon USMB treatment.

**Figure 6.**
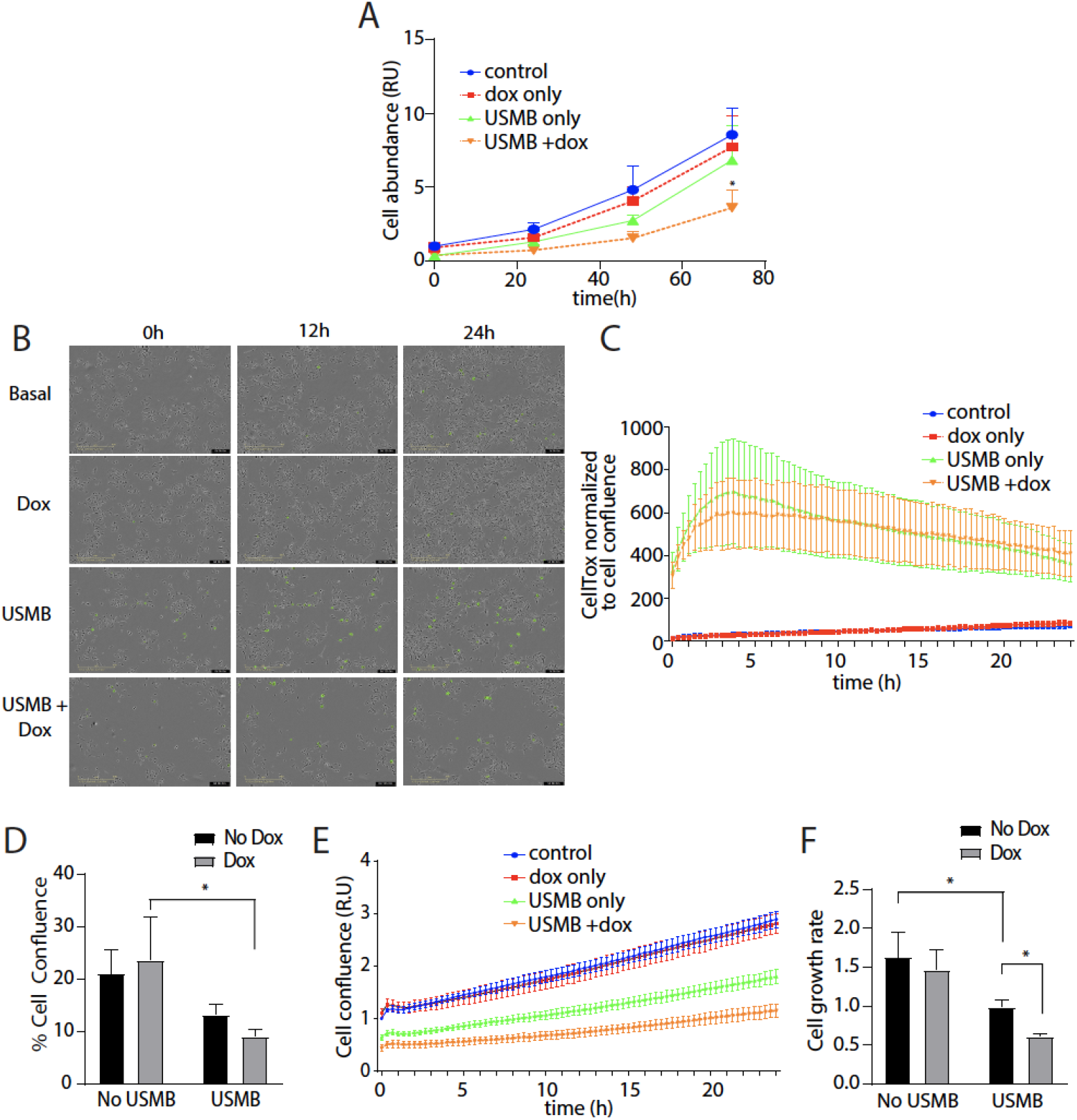
AMPK is required for MDA-MB-231 cell growth following USMB treatment. (A) MDA-shAMPK cells were treated ± 3μM doxycycline for 48 h (dox) followed by USMB treatment as indicated. Cell counting assay was performed every 24 h; measurements were normalized to the untreated control condition and shown as mean ± SD (n=5). Statistical analysis was performed by two-way ANOVA with Tukey’s multiple comparison test. *, p < 0.05, relative to corresponding timepoint in control (no doxycycline, no USMB treatment) condition. (B) Representative images from time-lapse recording of MDA-shAMPK cells treated with a solution containing CellTox. Representative images shown as overlay of brightfield and green fluorescence. Scale bar, 400μm. (C) Quantification of CellTox positive objects (cells) normalized to cell confluence shown mean ± SEM of n=3 independent experiments. (D-F) Quantification of Incucyte measured phase confluence of MDA-shAMPK cells treated as indicated with 3μM doxycycline for 48 h to silence AMPK followed by USMB treatment. Shown in (D) is the percent confluence at t=0 min following USMB, normalized to untreated control measured ± SD (n=3). Statistical analysis was performed by two-way ANOVA with Sidak’s multiple comparison. *, p<0.05. Shown in (E) is the relative confluence at regular intervals over 24 h following USMB treatment, normalized to untreated control ± SD (n=3). Shown in (F) is the result of determination of the rate of cell growth (slope of values of each individual experiment in E) shown as mean ± SD (n=3). Statistical analysis was performed by two-way ANOVA with Sidak’s multiple comparison. *, p < 0.05.

To further explore the role of AMPK in adaptation to USMB-induced stress, we simultaneously monitored MDA-MB-231 cell viability and proliferation using an automated environmentally-controlled microscope system (**Figure 6B** and Figure S1). Cell death was detected in real-time by labeling cells with CellTox Green and by normalizing against confluence to account for possible differences in cell abundance when assessing the fraction of non-viable cells. Consistent with past studies (Duan et al., 2021; Kun et al., 2009; Zhong et al., 2011), we found that MDA-MB-231 cells treated with USMB exhibited an immediate increase in CellTox Green positive cells in comparison to non-USMB exposed cells, indicating loss of cell viability (**Figure 6B, C** orange and green lines). Following USMB, MDA-shAMPK cells displayed a similar level of acute loss of cell viability by this method, whether or not they were treated with doxycycline to deplete AMPK (**Figure 6D**). These observations indicate that MDA-MB-231 cells do not rely on AMPK to enhance cell viability following USMB either acutely, as seen immediately (<5 min) after USMB treatment (**Figure 6D**), as well as over a 24h period following USMB (**Figure 6C**).

We next examined the long-term effects of USMB treatment, analogously to experiments in **Figure 6A**. MDA-shAMPK cells treated or not with doxycycline started at similar confluence (20%) and increased at similar rates over 24 h (**Figure 6E–F**). As expected, given the immediate, acute reduction in cell viability, USMB treatment significantly reduced initial confluence, which was similar whether or not cells were treated with doxycycline to suppress AMPK expression (**Figure 6D–E**). However, over 24 h, the increase in confluence upon USMB treatment was attenuated by AMPK depletion (**Figure 6E–F**).This indicates that in MDA-MB-231 cells, AMPK is essential for the long-term recovery from USMB treatment, and that the loss of AMPK manifests as a reduction in cell proliferation following USMB.

To examine if AMPK could play a role in cellular adaptation after USMB treatment in other cell types, we examined cell viability and cell confluence in SUM149PT-shAMPK cells following treatment with USMB. First, as seen by a large increase in CellTox-positive cells, we found that USMB resulted in an immediate decrease in cell viability, irrespective of whether cells were or were not treated with doxycycline to suppress AMPK expression (**Figure 7 A, B).** Importantly, in the absence of doxycycline, and thus in cells expressing AMPK, SUM149PT-shAMPK cells showed a gradual recovery after USMB treatment, as seen by a reduction in the fraction of non-viable cells over time (**Figure 7B**, green line). In contrast, SUM149PT-shAMPK cells treated with doxycycline to suppress AMPK expression maintained an elevated level of cell death for at least 24 h following USMB (**Figure 7B**, orange line). SUM149PT-shAMPK cells suffered a cell proliferation defect after USMB that was seemingly less dependent on AMPK expression (**Figure 7C–D**). Together, these observations indicate that AMPK plays a critical role in cellular adaptation and recovery from USMB-induced stress in MDA-MB-231 and SUM149PT cells, though specific aspects of these effects may be cell type specific.

**Figure 7.**
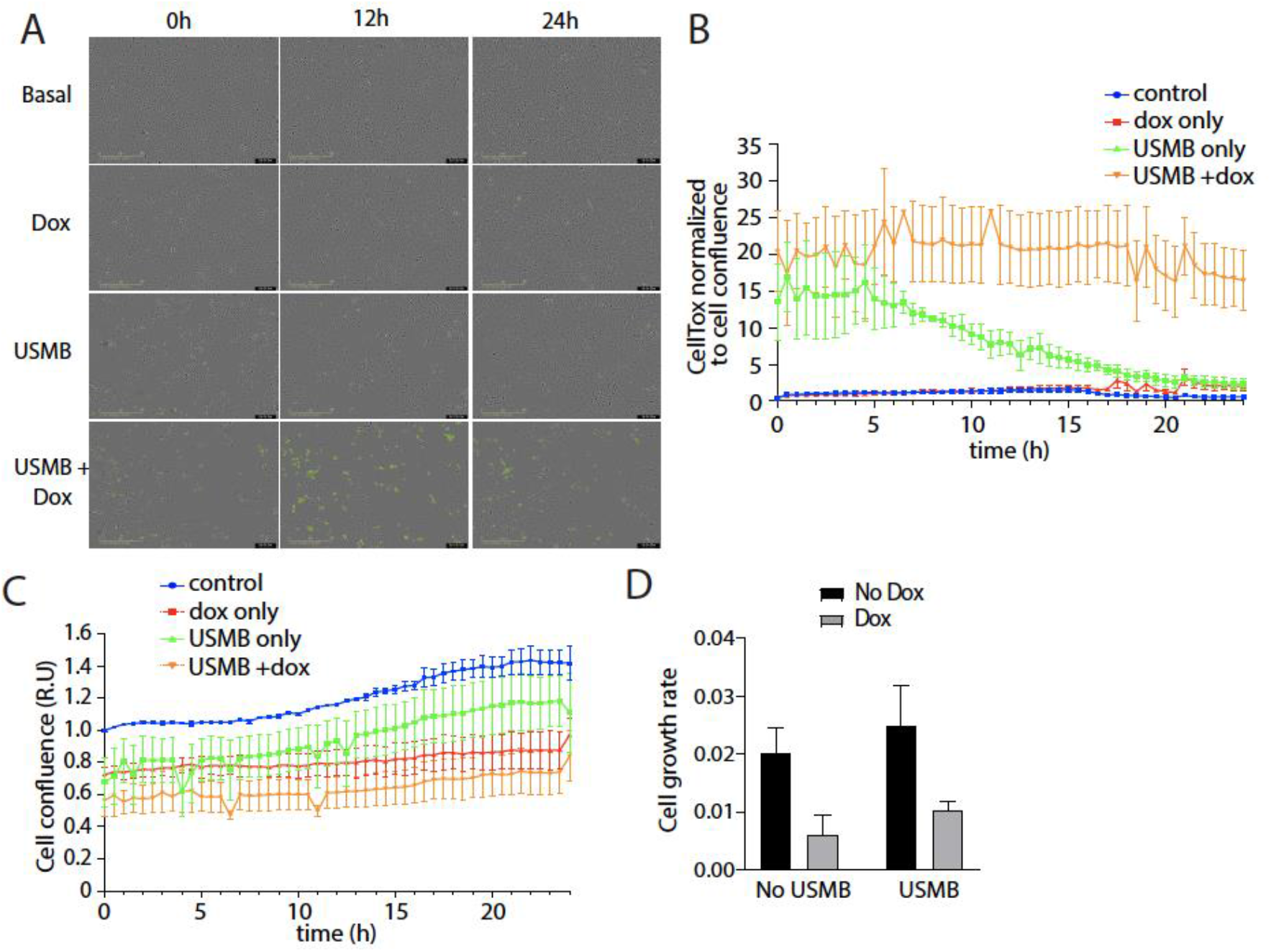
AMPK is required for SUM149PT cell survival following USMB treatment. SUM149PT-shAMPK cells were treated ± 3μM doxycycline for 48 h (dox) followed by USMB treatment as indicated. (A) Representative images from time-lapse recording of SUM149PT-shAMPK cells treated with a solution containing CellTox. Representative images shown as overlay of brightfield and green fluorescence. Scale bar, 400 μm (B) Quantification of total CellTox positive objects (cells) relative to the cell confluence, shown as mean ± SEM of n=3 independent experiments. Statistical analysis was performed by two-way ANOVA followed by Tukey post-test. All measurements of cell viability for the USMB+dox condition from 7-24h after USMB are significantly different (p < 0.05) compared to USMB only treated cells. (C) Quantification of phase confluence of SUM149PT-shAMPK cells treated as indicated with 3 μM doxycycline for 48 h to silence AMPK followed by USMB treatment. Shown is the relative confluence at regular intervals over 24 h following USMB treatment, normalized to untreated control, shown as the mean ± SEM of n=3 independent experiments. (D) Rate of cell growth (slope of values of each individual experiment in C) shown as the mean ± SEM of n=3 independent experiments.

## Discussion

The potential theragnostic applications of ultrasound in combination with microbubbles for cancer is gaining interest as unique cell biological effects of USMB are revealed and characterized. Until now, USMB treatment has been well-established to facilitate drug delivery into cells through sonoporation and enhanced endocytosis mechanisms (Misra et al., 2021). The focus of USMB for drug delivery has been largely centered on bringing mostly impermeable extracellular materials (drugs or nucleic acids) into the cell through sonoporation as opposed to the cell biological effects associated with loss of intracellular material such as its use to specifically kill target cancer cells. Here, we show that USMB reduces the levels of many intracellular metabolites, in particular amino acids, and glycolytic and citric acid cycle intermediates. To recover from this insult, MDA-MB-231 and SUM149PT cells rely on AMPK activation. Thus, inhibiting AMPK during USMB treatment may represent a strategy to enhance killing of cancer cells in a tumor.

Our metabolome analysis revealed that USMB treatment causes a reduction in the intracellular levels of a broad range of metabolites. A possible mechanism for this reduction in metabolite levels is the exchange of cytosolic materials from the cell interior to the extracellular milieu. However, the direct loss of metabolites by diffusion seems unlikely to account for all reductions in metabolite levels, such as those of the TCA cycle that should be found within mitochondrial matrix, and thus are less likely to directly efflux from the cell as a result of USMB-induced plasma membrane pores, suggesting instead that anaplerosis and/or cataplerosis contribute to changes in some metabolites following USMB treatment. While beyond the scope of the current study, future work that may delineate the precise molecular mechanisms underpinning USMB-induced changes in metabolic flux may further reveal the metabolic impact of USMB on cancer cells.

USMB treatment leads to a reduction in ATP levels within the cell, and AMPK is required for restoring cellular ATP within 2 h following USMB treatment (**Figure 4**). There are several mechanisms by which AMPK may promote metabolic adaptation resulting in replenishing of ATP levels (Herzig and Shaw, 2018; Rahmani et al., 2019). For instance, AMPK activation increases glucose uptake and promotes glycolysis, by acutely increasing the transport of glucose via facilitative glucose transporters such as GLUT1 (Wu et al., 2013) and phosphorylation of phosphofructokinase 2 (Marsin et al., 2000; Wu and Wei, 2012) as well as by transcriptional regulation of various glycolytic enzymes (Faubert et al., 2013; Marsin et al., 2000; Wu and Wei, 2012). Furthermore, ACC activity is suppressed as a result of its phosphorylation by AMPK and its activity can be further abated by reduced citrate concentrations (Hunkeler et al., 2018). Hence, ACC inactivation by AMPK suppresses fatty acid synthesis while increasing fatty acid oxidation (Hardie and Pan, 2002), leading to restoration of ATP levels (Hardie and Pan, 2002). AMPK activation also represses additional metabolic pathways such as hexosamine (Scott and Oakhill, 2017; Zibrova et al., 2017) and mevalonate biosynthesis (Saha et al., 2018). Future work should strive to define which of these mechanisms primarily drives cellular adaptation to USMB-induced loss of metabolites and ATP.

Our results further indicate that AMPK activation supports cancer cell proliferation. While many studies report that activation of AMPK, such as with biguanides like metformin, suppresses proliferation of cancer cells (Cao et al., 2019; Faubert et al., 2015; Hardie, 2015), other studies have shown that in certain contexts, AMPK activation is required to support cell proliferation. In lung cancer driven by oncogenic KRAS mutation, abrogation of AMPK led to substantial impairment of tumor growth, revealing the role of AMPK to promote lysosome expansion and autophagy (Eichner et al., 2019). The tumor-promoting functions of AMPK are also associated with AMPK enhancement of glucose transporter expression and glucose transport (Wu et al., 2013), enhanced carbohydrate metabolic capacity supported by heightened glycolytic enzyme expression (Bando et al., 2005; Herzig and Shaw, 2017; Marsin et al., 2000), and improved mitochondrial capacity, such as that resulting from AMPK regulation of PGC1α (Herzig and Shaw, 2017; Jäer et al., 2007). Overall, AMPK is important for at least some cancer cells to adapt following USMB treatment, thus supporting cell proliferation.

### Limitations of study

The sonoporation phenomenon has been extensively studied in a number of models; *in vivo* (animals, solid tumors) (Kotopoulis et al., 2014; Todorova et al., 2013) and *in vitro* (cultured cancer cells and non-cancer cells) (Haugse et al., 2019; Qu et al., 2018; Wang et al., 2018). While we have shown activation of AMPK following USMB treatment in three TNBC cell types (MBA-MB-231, SUM149PT and BT20), other cell types and cell lines may respond and behave differently when exposed to USMB (Qu et al., 2018). Moreover, the nature and extent of the cell biological effects of USMB may depend on various parameters of ultrasound settings and microbubble properties, such as concentration and formulation of microbubbles, and ultrasound parameters (soundwave, microbubble formulation, duration of treatment), which were previously defined for the cell lines we examined here.

## Acknowledgements

We thank Dr. Ivan Topisirovic (Lady Davis Institute and McGill University, Montreal) for helpful discussions and critical reading of this manuscript. This work was supported by Discovery Grants from the Natural Sciences and Engineering Research Council to C.N.A. (RGPIN-2016-04371) and R.K. (RGPIN-201505941) and by an Engage Grant from the Natural Sciences and Engineering Research Council, the Canada Research Chair Program (950-232333), the Early Researcher Award (ER13-09-042) from the Ministry of Research and Innovation, Government of Ontario, the Canada Foundation for Innovation (32957) and respective contributions from Ryerson University to R.J.B.

## Author Contributions

Conceptualization: L.L., R.J.B., C.N.A., R.K., Investigation and formal analysis: L.L., O.U., writing – original draft: L.L., R.J.B., C.N.A., R.K., writing – review & editing – O.U., supervision and funding acquisition: R.J.B., C.N.A., R.K.

## Supplemental Information

### Methods

#### Materials

DEFINITY microbubbles were obtained from Lantheus Medical Imaging Inc (Saint-Laurent, QC).

Antibodies used for immunofluorescence microscopy were as follows (with dilutions used also indicated): Rabbit mAb anti-phospho-AMPK rabbit mAb anti-phospho-Acetyl-CoA Carboxylase (Ser79, cat#11818, 1:400), rabbit mAb anti-Acetyl-CoA Carboxylase (cat#3662, 1:1000) from Cell Signaling Technology (Danvers, MA), goat anti-rabbit Alexa Fluor® 488 and donkey anti-rabbit Alexa Fluor® 647 from Jackson ImmunoResearch Laboratories (West Grove, PA).

Antibodies used for Western Blot were as follows (with dilutions used also indicated): anti-GAPDH rabbit mAb (cat#2118, 1:1000), anti-Phospho-Acetyl-CoA Carboxylase rabbit mAb (Ser79, cat#11818, 1:1000), anti-Acetyl-CoA Carboxylase rabbit pAb (cat#3676, 1:1000), anti-AMPKα rabbit pAb (cat#2532, 1:1000), anti-Clathrin heavy chain rabbit mAb (cat#4796 1:1000) were obtained from Cell Signaling.

Dorsomorphin (Compound C) was obtained from Abcam (Cambridge, MA). Digitonin was obtained from Promega (Madison, WI, cat#G9441, 1:1000). Lucifer Yellow potassium salt was obtained from ThermoFisher (cat#L1177). Doxycycline hydrochloride was obtained from BioBasic (Markham, ON, cat#DB0889). CellTox Green cytotoxicity assay was obtained from Promega (cat#G8741).

#### Cell Culture and siRNA-mediated gene silencing

The human breast cancer cell line MDA-MB-231 was obtained from American Type Culture Collection, ATCC (HTB-26). MDA-MB-231 cells were cultured in RPMI 1640 media supplemented with 10% fetal bovine serum (FBS, ThermoFisher Scientific) and 5% penicillin/streptomycin (ThermoFisher Scientific) in a 37°C and 5% CO2 controlled by an incubator system (ThermoFisher Scientific).

The human breast cancer cell line SUM149PT was a kind gift from Dr. Daniel Durocher (Lunenfeld-Tanenbaum Research Institute, Toronto, Canada). SUM149PT cells were cultured in Ham’s F-12 media (Gibco) supplemented with 5% fetal bovine serum (FBS, Gibco), 10mM HEPES (Sigma-Aldrich), 1ug/mL hydrocortisone (Sigma-Aldrich), 5μg/mL insulin (Sigma-Aldrich), and 5% penicillin/streptomycin (ThermoFisher Scientific) in a 37°C and 5% CO2 controlled by an incubator system (ThermoFisher Scientific).

The human breast cancer cell line BT-20 was obtained from American Type Culture Collection, ATCC (HTB-19). BT-201 cells were cultured in Minimal Essential Media Earle’s Eagle media supplemented with 10% fetal bovine serum (FBS, ThermoFisher Scientific) and 5% penicillin/streptomycin (ThermoFisher Scientific) in a 37°C and 5% CO_2_ controlled by an incubator system (ThermoFisher Scientific).

siRNA silencing of AMPK was performed as previously described (Ross et al., 2015). Custom synthesized siRNA sequences were designed to silence expression of both AMPK α1/2 isoforms (**Table S1**). MDA-MB-231 cells were grown to 40% confluency before being subjected to a single round of siRNA transfection using Lipofectamine RNAiMAX transfection (ThermoFisher Scientific, cat#13778075) performed according to manufacturer’s instructions. Cells were exposed to the transfection mixture for 6 h, allowed to rest for 48 h in fresh culture media before treatment.

#### Generation of MDA-MB-231-shAMPK and SUM149PT-shAMPK Stable Cell Lines

To generate stable MDA-MB-231 cells and SUM149PT cells that conditionally silenced AMPK, we designed a sleeping beauty transposon system containing a stable expression of a BFP-reporter gene and a doxycycline inducible gene (Kowarz et al., 2015) to encode expression of a short hairpin sequence that can target both isoforms of the catalytic subunit (α1, α2) AMPK mRNA (**Table S1**), and a puromycin resistance gene for cell selection was used.

To generate the pSBtet-shAMPK-BP construct, two initial constructs were initially obtained. MSCV p2GM AMPK alpha2hp1 alpha 1hp1 was a gift from Russell Jones (Addgene plasmid #89492) (Faubert et al., 2013). pSBtet-BP was a gift from Eric Kowarz (Addgene plasmid # 60496; http://n2t.net/addgene:60496; RRID:Addgene_60496) (Kowarz et al., 2015). pCMV(CAT)T7-SB100 was a gift from Zsuzsanna Izsvak (Addgene plasmid # 34879; http://n2t.net/addgene:34879; RRID:Addgene_34879) (Mátés et al., 2009). The shRNA scaffold and AMPK shRNA sequence from MSCV p2GM AMPK alpha2hp1 alpha 1hp1 (Table S1) was cloned into pSBtet-BP to replace the luciferase gene (construct generated by Biobasic, Markham, ON, Canada) to generate pSBtet-shAMPK-BP.

MDA-MB-231 cells were cultured to confluence before transfection with FuGENE Transfection Reagent (Promega, E2311) of pSBtet-shAMPK-BP and pCMV(CAT)T7-SB100. Following transfection, cells were washed with PBS and allowed to recover for 24 h in culture medium. Cells were then dissociated and cultured in a new dish for 2 weeks in culture medium containing 2 μg/mL puromycin. Selected colonies were pooled and FACS sorted based on blue fluorescent protein (BFP) expression (expressed constitutively by the original pSBtet-BP and newly generated pSBtet-shAMPK-BP plasmids). Using FACS, the cells with the greatest BFP intensity (top 30%) of the population was selected and subsequently cultured in 2mg/mL puromycin containing RPMI media.

#### Microbubbles

DEFINITY microbubbles (Lantheus Medical Imaging Inc., Saint-Laurent, QC) were activated using a Vialmix for 45 seconds according to the manufacturer’s protocol. Microbubbles were used within 1 hour of activation. A 180 μL volume of activated microbubbles was diluted in 570 μL of room temperature PBS to a total volume of 750 μL. The mixture was resuspended by hand agitation and injected into the treatment vessel immediately prior to US exposure, giving a concentration of 10μL of microbubbles per mL of media.

#### Ultrasound Treatment

The study design we used is outlined in Figure 1, and as previously described (Fekri et al., 2019). Briefly, a six-well plate with adherent cells was filled with 16 mL of media and then exposed to an ultrasound wave at 500 kHz pulse centre frequency. We used a single element flat transducer with 32 mm element diameter focused at 85 mm and a −6 dB beam width of 31 mm at the focal point (IL0509GP, Valpey-Fisher Inc. Hopkinton, MA, USA), 690 kPa peak negative pressure, 32 μs pulse duration (16 cycle tone burst) at 1 kHz pulse repetition frequency corresponding to 3.2% duty cycle, for 60 seconds in the presence of prepared microbubbles. An arbitrary waveform generator connected to a power amplifier (AG series Amplifier, T&C power conversion Inc., NY) transmitted ultrasound parameters to the ultrasound transducer.

#### Immunofluorescence Staining

Immunofluorescence staining was performed as previously described (Bautista et al., 2018; Fekri et al., 2019). Cells were seeded onto coverslips in growth medium. Following treatment, cells were washed 2X with PBS (with Mg^2+^ and Ca^2+^) and then immediately placed on ice and fixed with 4% paraformaldehyde for 10 min, quenched with glycine for 5 min, and permeabilized with 0.1% triton X-100 and glycine solution for 5 min. Cell samples were subject to blocking using SuperBlock (ThermoFisher Scientific, cat#37515) according to the manufacturer’s protocol. Cells were stained with anti-Phospho-Acetyl-CoA Carboxylase (Cell Signaling. #11818) using a 1:400 dilution in a PBS supplemented with 1% BSA (1% BSA-PBS) for 1 h at room temperature by the inverted drop technique. Cells were washed over ten times with PBS, then stained with AffiniPure Goat anti-rabbit Alexa Fluor488-conjugated antibody (1:1000 dilution in 1% BSA-PBS) (Jackson ImmunoResearch Laboratories Inc. #111-545-144) for 1 h at room temperature in the dark. Cells were washed over ten times with PBS and mounted onto coverslips with Dako Fluorescent Mounting Medium (Agilent Technology #S3023). The slides were incubated at room temperature overnight to solidify before being stored in −20°C storage until visualization.

#### Epifluorescence Microscopy and Image Analysis

All samples were visualized on an inverted microscope by widefield epifluorescence microscopy (Olympus IX83 epifluorescence microscope, Hamamatsu ORCA FLASH4.0 C11440-22CU camera, run by Olympus cellSens software). For each experiment, randomly chosen fields were selected for acquisition of pS79-ACC with a 40x magnification using 100ms exposure time. Using ImageJ software, total fluorescence signal was individually measured for >100 cells with background correction. Fluorescent intensity was normalized to untreated control and expressed as fold change. Ordinary one-way ANOVA with Tukey’s multiple comparisons test was performed for the conditions.

#### Cell Lysis and Western Blots

Western blotting was performed as previously described (Bone et al., 2017; Fekri et al., 2019). Cells were washed twice with cold PBS and lysed with a cell scraper in Laemmli sample buffer supplemented with protease inhibitors (cOmplete™ EDTA-free Protease Inhibitor Cocktail, Roche, cat#1187358001), sodium orthovanadate (1mM), okadaic acid (1μM), bromophenol blue (0.004%), and 2-mercaptoethanol (10%). The protein lysates were heated to 65°C and passed through a 27.5 G needle. The lysates were separated on mini-PROTEAN TGX precast gels (Bio-Rad), and transferred onto FluroTrans PVDF (cat#PVM020C-099, PALL Life Sciences) using wet transfer method. The immunoreactive bands were developed using Luminata Crescendo Western HRP substrate (cat#WBLUR0100, Millipore Sigma) and detected using a ChemiDoc Touch imaging system (Bio-Rad).

#### GC-MS for steady state metabolite analysis

Metabolite analysis was performed as previously described (Hulea et al., 2018). Cells were counted via automated cell counting method and 4.50×10^5^ cells were seeded in triplicates (n=3) in 6-well plates 24 hours prior to treatment. Treated cells were placed on ice and washed twice with ice cold saline solution (0.9% NaCl). Subsequently, the cells were scraped from the plate while placed on dry ice in 300μL of 80% methanol chilled to −20 °C. The extracts were collected and transferred to prechilled tubes. To ensure complete collection of the material, 300 μL more of 80% methanol was added to the leftover cells in the wells, collected and added to the previously collected 300 μL fraction. Cell extracts were sonicated and the supernatants were extracted via centrifugation. After the supernatants were transferred into Eppendorf tubes, 750 ng of myristic acid-D27 was added to each sample to serve as an internal standard. The supernatants were dried overnight via CentriVap at 4°C (Labconco). The dried residues were dissolved in 30 μL of pyridine containing methoxyamine-HCl (10 mg/mL) (MilliporeSigma). Samples were then incubated for 30 min at 70°C. Afterwards, 70 μL of N-tert-Butyldimethylsilyl-N-methyltrifluoroacetamide (MTBSTFA) was added to the sample mixtures and incubated for 1 h at 70 °C. 1 μL of each sample mixture was used for GC–MS analysis. The GC-MS machine and software were obtained from Agilent. Data analyses were performed on ChemStation and MassHunter software (Agilent). We quantified the peak area for each metabolite based on known ion peaks, and normalized this to sample protein content and to an internal loading standard Myristic D-27. Individual metabolite values were averaged and normalized to untreated controls to determine fold changes for each metabolite compared to untreated controls.

#### ATP Assay

Luminescent ATP detection assay kit (abcam, ab113849) was used to quantify ATP according to manufacturer’s protocol. Briefly, cells seeded in a six-well plate were lysed with 200μL of the provided detergent and equal volume of substrate solution was added before luminescence detection on a BioTek Synergy HTX Multi-Mode Reader. Results were normalized to the total protein content for the respective treatment as determined though the BCA assay (ThermoFisher, cat#23227). The results were expressed as fold change (means ± SD) over untreated controls. Ordinary one-way ANOVA followed by Tukey’s multiple comparisons test was performed.

#### Cell Counting Analysis

Treated cells were washed with PBS and then dissociated with 200μL Trypsin-EDTA (Thermofisher, cat#25200065). 200μL of complete culture media was added to end the trypsinization reaction. 10μL of cells was collected in a tube and mixed with 10μL of 0.4% trypan blue. Cells were counted with a Countess II FL (ThermoFisher Scientific). Three technical replicates for each condition were recorded. Results were averaged and normalized to untreated starting conditions. The results were expressed as fold change of cell number (mean ± SD) over initial untreated control. Two-way ANOVA followed by Tukey’s multiple comparisons test was performed.

#### Cytotoxic assay and cell confluency measurement using the Incucyte

Cell death was assessed using CellTox Green Cytotoxicity Assay reagent (Promega G8741). 1μL of the dye was diluted in 10mL of media, and this mixture was added to the cells 5 min after USMB treatment. Cytotoxicity and viability were measured using an Incucyte S3 Live-Cell Analysis System (Sartorius). Images were acquired at regular intervals under 10x magnification using brightfield phase contrast and 100ms exposure of 460nm excitation, 524nm emission for green fluorescence. Analysis was performed with Incucyte Plategraph for CellTox positive cells and phase confluence. Two-way ANOVA followed by Tukey’s multiple comparisons test was performed.

---

**Supplemental Table 1.**
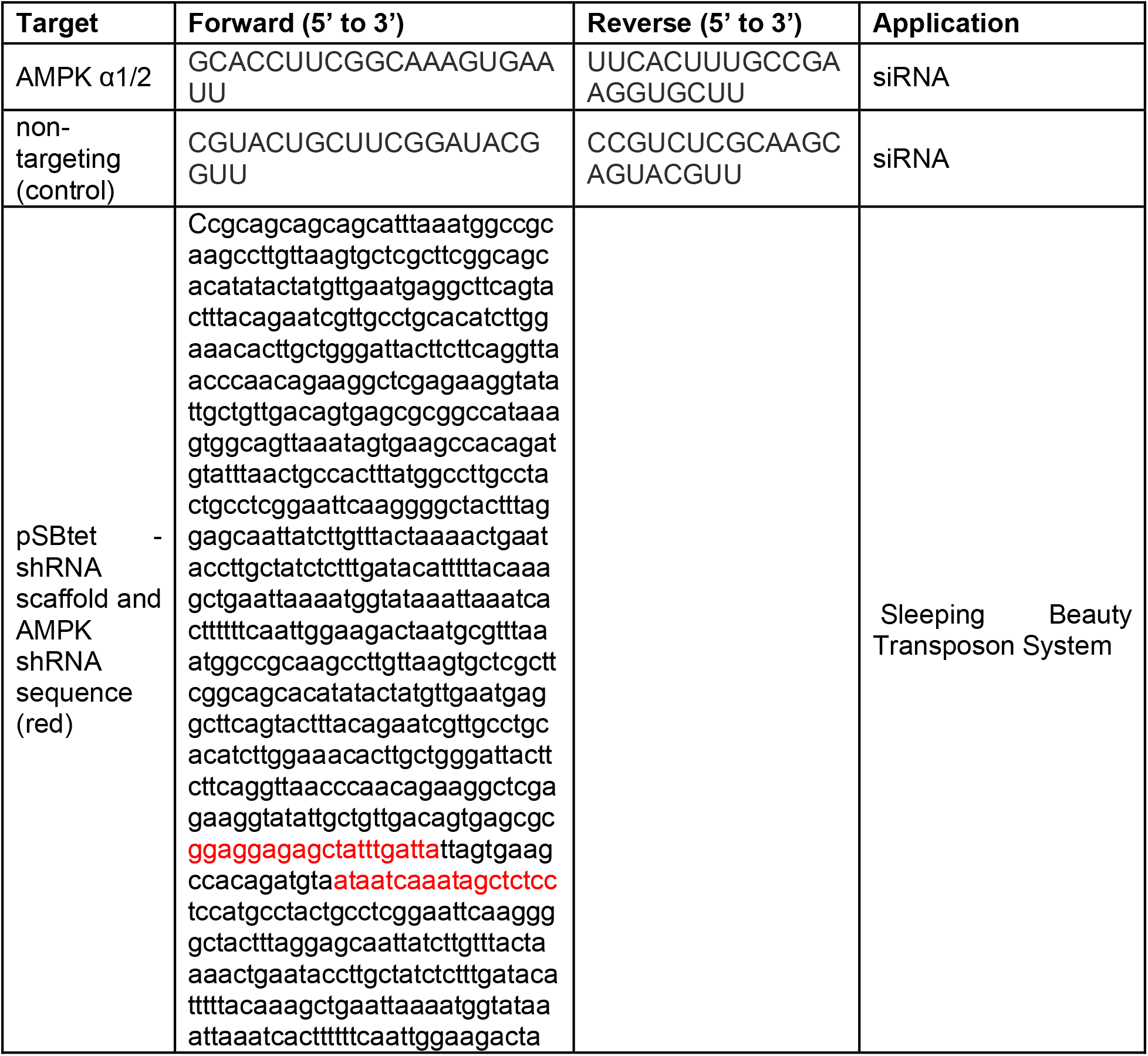
Sequences used for siRNA and shRNA. Shown in red is the AMPK targeting sequence within the shRNA scaffold.

**Supplemental Figure 1.** Time-lapse recordings of MDA-shAMPK cells treated with a solution containing CellTox Green, related to **Figure 6B**. MDA-shAMPK cells were treated ± 3μM doxycycline for 48 h (dox) followed by USMB treatment as indicated in Figure 6. (A) basal condition (no USMB or doxycycline) (B) doxycycline treated condition (AMPK silenced) (C) USMB only condition (no doxycycline) (D) doxycycline treated sample followed by USMB treatment.

